# Eavesdropping and audience effect in fighting fruit flies reveal feedback between social information use and production

**DOI:** 10.1101/2023.09.01.555942

**Authors:** Marina Hutchins, Julia Saltz

**Affiliations:** Rice University, Department of Biosciences, Houston (TX), USA

**Keywords:** aggression, audience effect, Drosophila melanogaster, eavesdropping, social information

## Abstract

Social information use and production occur simultaneously during social learning events; however, these two processes are often studied independently. We propose that social information use and production represent a feedback loop: eavesdroppers influence the behaviors of those being observed, which then alters the eavesdroppers’ social decisions. Here, we used *Drosophila melanogaster* to investigate the relationship between social information use and production in an aggressive context. We tested whether paired males altered their aggression when they were observed by (i) a male; (ii) a female, or; (iii) no one. Next, we allowed each individual eavesdropper to interact with the paired males that they just observed, and measured their social behaviors. In addition, we tested how naïve flies responded when interacting with paired males. We found that paired males reduced their aggression when observed by a male, representing a change in social information production. In response, male eavesdroppers were overall less aggressive towards the more aggressive of the paired males they had observed, but their response to social information was also modulated by their genotype. In contrast, female eavesdroppers did not elicit changes to the behavior of the paired males, nor did we find evidence that they used the social information collected to make mating decisions. Overall, we found support for the feedback loop in males, but not females: information was exchanged bi-directionally during eavesdropping events, resulting in the modification of behavior by all parties. Utilizing the social information feedback framework can contribute to a better understanding of how social behaviors and animal communication are expressed and evolve.

**Competing Interests:** None

**Significance Statement:** Animals often observe the social interactions that occur between others, but the effects of both watching and being watched on behavior are rarely studied together. Using a novel approach, we studied how an audience influenced and learned from an aggressive contest in fruit flies. A male audience caused contest participants to decrease their aggression, and the male who was watching later showed decreased aggression toward the contest winner. Surprisingly, we did not see any effect of a female audience on male-male aggression, and females’ later behavior was not altered by observing a contest. Together, these results show how observing a fight led to changes in both the behavior of the contest participants and the male audience, thus suppressing group levels of aggressive behavior. Accounting for this interaction between social information use and production is necessary to understand social behaviors.

## Introduction

Individuals are able to acquire social information by detecting information about the behavioral interactions of others through a variety of sensory modalities and mechanisms (Heyes, 1994; McGregor & Peake, 2000; Wagner & Danchin, 2010). We refer to these third-party conspecifics as ‘social eavesdroppers’: individuals who change their behavior after having access to information about the interactions of others. (Bonnie & Earley, 2007; Dall et al., 2005; Danchin et al., 2004; McGregor, 2005, Tibbets et al., 2020). Social eavesdropping could provide a low-risk alternative to direct interactions with others, as eavesdroppers can learn about the quality or ability of another individual without engaging in interactions that could be risky or costly (McGregor & Peake, 2000; Mery et al., 2009). For instance, females of many species have been shown to eavesdrop on the mate choice decisions of other females and then copy those preferences, an example of eavesdropping as a cost-avoidance strategy (Vakirtzis, 2011).

In many cases, an interacting pair can detect the presence and sometimes the characteristics of a social eavesdropper, which may lead to a shift in behavior known as the ‘audience effect’ (Fichtel & Manser, 2010; McGregor, 2005, Coppinger et al. 2017). The audience effect is widespread and well-documented in a variety of animal species, including fish (Doutrelant et al., 2001; Plath et al., 2008), birds (White et al., 2002), primates (Townsend & Zuberbuhler, 2009), and humans (Bond & Titus, 1983). For example, substantial work has demonstrated that in primates, the intensity and/or frequency of alarm calls performed by an individual depends on the characteristics of the audience members around them (Chapman et al., 1990; Pollick et al., 2005; Wheeler, 2008).

While social eavesdropping and the audience effect are two well-described phenomena, they have historically been studied as two distinct processes (McGregor, 2005). Yet, social eavesdropping and the audience effect can occur in tandem during social learning events (Bonnie & Earley, 2007; Chimento et al., 2022; Cruz & Oliveira, 2015; Earley & Dugatkin, 2002). During these events, the expression of social behaviors – and thus, the production of social information – may be altered depending on the presence and identity of the eavesdropping individual, i.e., via audience effects (Desjardins et al., 2012; Fitzsimmons & Bertram, 2013; Plath et al., 2008). At the same time, the social eavesdropper can use the social information by changing their own behavior depending on what they observed from the interacting pair (Chimento et al., 2022; Peake et al., 2001). Taken together, we can hypothesize a feedback loop between social information production and use, in which the presence of a social eavesdropper alters the social environment of the interacting pair, which in turn influences the behavior of both the interactants and the eavesdropper.

The existence of a feedback loop between the production and use of social information has the potential to alter our current understanding of how social behaviors are expressed and have evolved, especially in group contexts. If there is no feedback between these processes, then research dedicated to studying social information use independently from social information production would be sufficient to capture the key social dynamics that emerge from these two phenomena. However, if a feedback loop does exist, then studies focused on only one-half of the feedback loop at a time - i.e., studying interacting pairs without eavesdroppers or vice versa - ignore critical participants during these interactions. Under feedback loops, bi-directional information exchange between the social eavesdroppers and the observed pair has the potential to modify the behavior of all parties, thus influencing our predictions about social learning and animal communication in social groups. For instance, early work investigating social information use in female cowbirds initially found that rearing environment did not influence females’ preference for male song between two cowbird subspecies (King and West 1983). However, researchers later discovered that females modified the social environment that they were reared in –an unaccounted for audience effect that altered their interpretation of their findings (King & West 1987). Indeed, recent theory has highlighted the need to investigate how variation in social information use and production can influence decision-making by incorporating theory and data about both audience effect and social eavesdropping simultaneously (Chimento et al., 2022; Turner et al., 2023).

In animal societies, aggression is an important social signaling behavior that influences social relationships, foraging, and competition (King, 1973; Duque-Wilckens et al., 2019). Aggression often takes place in an environment where social eavesdroppers are present and thus may be able to observe contests between others and extract information about the interactants’ fighting ability (Oliveira et al., 1998; Peake et al., 2001). Previous work in a variety of organisms has shown that social eavesdroppers can obtain information from the aggressive behaviors performed during a fight, such as threat displays or overt physical interactions, or the contest outcome and dynamics, such as the identity of the winner and loser (Johnstone, 2001; Magnhagen, 2006; Oliveira et al., 1998; Tibbetts et al., 2020; Van Breukelen & Draud, 2005).

Decades of research, primarily in fish, has been dedicated to investigating how individuals change their aggressive behavior in the presence of an audience, or how the outcome of a contest affects the future behavior of social eavesdroppers(see Peake & Mcgregor, 2004 for review). However, there are currently scant empirical studies identifying feedbacks, if any, between social information production and use in an aggressive context. For example, many studies of social eavesdropping are conducted by allowing an individual to observe a fight between others, while preventing the eavesdropper from being detected by the demonstrators (e.g., using one-way mirrors; Magnhagen, 2006; Oliveira et al., 1998; Peake et al., 2006). Making the eavesdropper undetectable to the interacting pair eliminates any possible audience effects; however, in many circumstances, the interacting individuals often can detect and respond to the presence of the eavesdropper, and *vice versa* (Zuberbühler, 2008). Thus, these experiments are not fully capturing and accounting for the information transfer that happens in social groups in the wild during social eavesdropping events.

Here, we utilized the social information feedback framework to investigate social decision-making in the context of aggressive interactions. To investigate the proposed feedback loop between social information production and use, we combined standard methods used for testing how individuals produce and how individuals use social information into a single experiment. In the first phase of each trial, we allowed individuals to eavesdrop on two males engaged in an aggressive contest – this observation phase allowed us to test for variation in how the paired males produced social information. In the second phase of the trial, the social eavesdroppers could then interact with both aggressors directly, which allowed us to test how the produced social information influenced the eavesdropper’s behavior. Measuring both social information use and production in our experimental trials allowed us to directly examine how an individual’s ability to influence the production of social information results in differences in social information use. We repeated this approach with 3 different genotypes of social eavesdroppers, which allowed us to test for differences in access to social information production and the robustness of the effect of social information use across different genotypes.

We predict that the presence, sex, and genotype of an eavesdropper will drive changes in the behavior of the interacting aggressive pair (Fitzsimmons & Bertram, 2013; Matos & McGregor, 2002). We further hypothesize that the social information observed by the eavesdroppers will alter their behaviors (i.e., aggression in males, or mating in females), towards one or more of the paired males (Aquiloni et al., 2008; Chan et al., 2008; Magnhagen, 2006). Observing both of these results would support the existence of a feedback loop between these processes. Further, any of these processes may be modified by the genotype of the eavesdropper.

## Methods

### Study system

*Drosophila melanogaster*, the fruit fly, is an excellent study system for investigating the relationships between social information use and production because of their well-characterized suite of behavioral interactions and amenability to rigorous laboratory experiments. D.melanogaster forms social groups on their food sources both in nature and in the lab, and social interactions, such as aggression, take place in an environment with ample conspecific social eavesdroppers (Saltz, 2013; Stamps et al., 2005; Wertheim et al., 2006). Fruit flies have a well-studied repertoire of aggressive behaviors (Chen et al., 2002; Saltz, 2013), and the outcomes of aggressive contests can influence later behavior (Trannoy et al., 2016). Fruit flies also have the necessary cognitive capacity for social information use (Danchin et al., 2018; Mery et al., 2009; Nöbel et al., 2018).

### Genotypes and Rearing

We used inbred fly lines that were originally derived from a wild population in Raleigh, NC (MacKay et al., 2012). We used 3 different genotypes of social eavesdroppers: 136, 229, and 429 (note that genotypes are “named” using numbers, and their numbers are arbitrary and have no numeric meaning). We standardized the genotypes of the paired males to ensure that we could directly link any effect of eavesdropper sex and presence to the interacting pair’s social information production. For the standardized paired males, we used one male from genotype 365 and one male from genotype 765. (Again, genotype numbers are arbitrary and have no further significance). Flies for this study were reared in vials under standard conditions that minimized variation in larval density: 10 females and 10 males were placed into a vial containing standard fly food medium. Unmated progeny from these vials were then collected the morning that they eclosed, and subsequently housed in individual vials to prevent any social experiences. Flies were aged for 3 or 4 days before experimental trials and were maintained on a 12:12 hour light:dark cycle.

### Behavioral Arena

We utilized a novel arena design to test for variation in social information use and production (Figure 1). The main part of the arena was composed of two Petri dishes (3 cm x 1 cm) taped together, with a small food patch (1.5 cm x 1 cm) placed on the bottom of the arena, which contained standard fly food medium and a dot of active yeast. There were two small holes in the arena: one on the side of the arena, for the insertion of flies via aspiration, and another hole in the center of the top of the arena. A pipette tip with a mesh base was placed over this hole, creating an observation tower for the eavesdropper fly to watch aggressive contests. This arena design mimics the perching behaviors of flies in natural conditions (Stamps et al., 2005). The mesh barrier that separated the eavesdropper fly from the paired males allowed visual, olfactory, and/or auditory information to be extracted from the aggressive contest (Versteven et al., 2017; Wang & Anderson, 2010). The location of the observation tower was directly over the food patch, where most aggressive interactions occur (Hoffmann & Cacoyianni, 1990). We observed occasional interactions at this mesh between the eavesdropper fly and the interacting pair, however, we did not observe any aggressive or courtship behaviors between the eavesdroppers and any member of the interacting pair.

**Figure 1.**
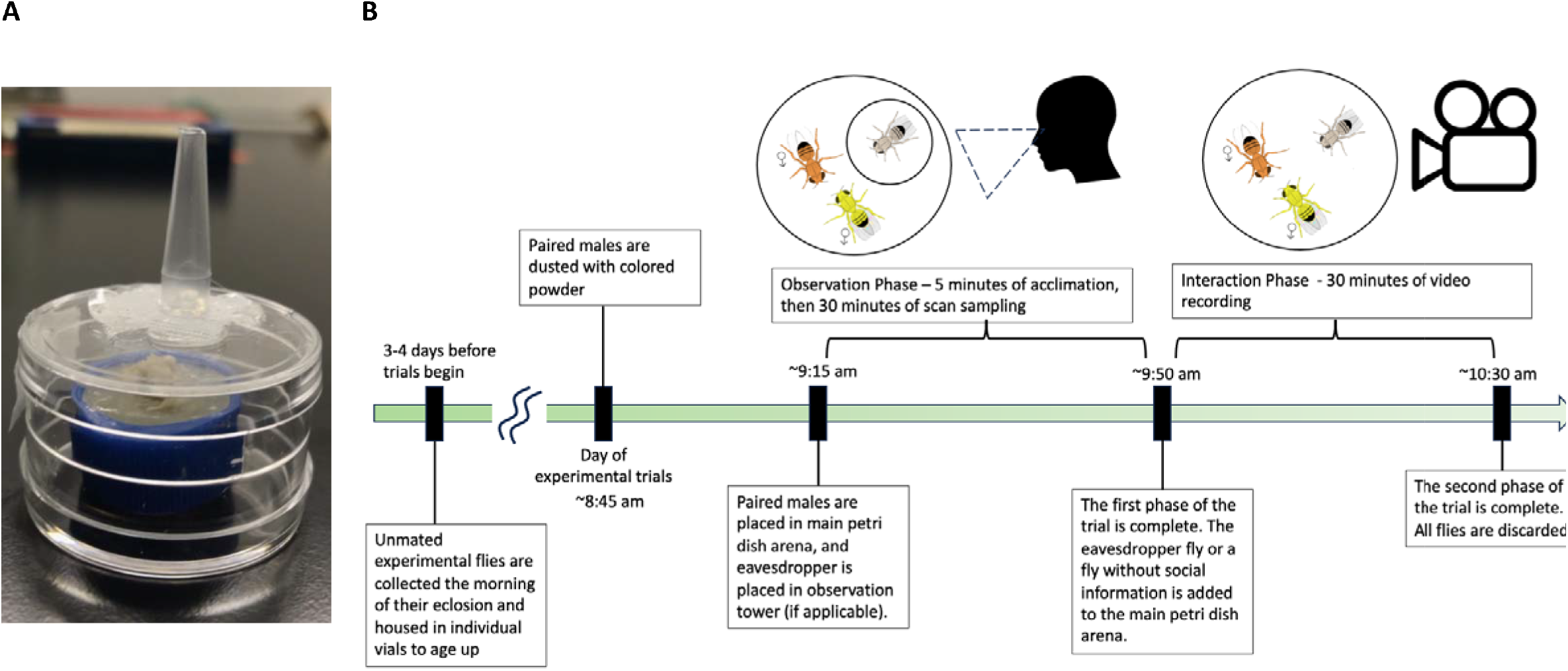
(A) The behavioral arena used to conduct experimental trials. The pipette tip on top housed an eavesdropper fly; the opening was then covered to prevent them from escaping. The paired males were inserted into the petri dish arena with the blue cap that was filled with standard food and topped with yeast paste. (B) Timeline overview of protocol for an experimental trial.

### Overview of trials

All flies utilized for the trials had no prior social experience, as they were socially isolated from the time of their eclosion until trials were conducted (see Genotypes and Rearing). Each trial consisted of two phases: the observation phase and the interaction phase. In the observation phase, an eavesdropper fly was placed into the observation tower while the paired males in the main part of the arena engaged in an aggressive contest. In the interaction phase, the same individual eavesdropper fly was added to the arena with the paired males they had just observed, and their social interactions with the males were recorded (Figure 1). After each trial was completed, all flies were discarded. Thus, each individual fly (eavesdropper or member of the interacting pair) was used in both phases of a single trial, and not reused in any other trials.

### Experimental Protocol

We randomly dusted the paired males with colored powder (orange or yellow) to create two distinct male phenotypes. This allowed human observers to easily differentiate between the two flies for scoring aggressive behavior, and this distinctive coloring also may have aided the eavesdropper flies in distinguishing between the two males (E. Danchin et al., 2018; Mery et al., 2009; Monier et al., 2018).We did not find any effect of color on male aggression or female mate choice (see Model Covariates in the Supplemental Material). The paired males were inserted into the main petri dish arena via aspiration, and the eavesdropper fly was placed into the observation tower. Flies were allowed to acclimate for five minutes before the observation phase of the experimental trials began.

During the observation phase of the experiment, we measured aggressive behaviors performed by the paired males for 30 minutes. We recorded five measures of aggressive behavior: lunging, fencing (both low- and high-level fencing), offensive wing threats, chasing, and boxing, as defined in Chen et al., 2002. Every two minutes, a human observer recorded all aggressive behaviors performed by the two male flies in the arena for 20 seconds. During this observation phase, the eavesdropper fly was able to observe the aggressive contest without interacting directly with the two males.

Social eavesdroppers rarely have the opportunity to make decisions about who to interact with (but see studies on mate choice in an aggressive context, Van Breukelen & Draud, 2005). Rather, experimental designs frequently designate the eavesdropping individuals to be randomly paired with only one of the contest participants (Earley & Dugatkin, 2002; Johnsson & Åkerman, 1998; Oliveira et al., 1998; Tibbetts et al., 2020). These designs prevent us from gaining insight into how individuals can process and utilize social information to make decisions about how they will behave toward other group members. Recent calls for the need to explain and understand social decision-making, especially in conflict settings, can be solved by utilizing an experimental design that gives the eavesdropper the choice of interacting with both contest participants (Hobson, 2020). Thus, immediately following the observation phase, we began the interaction phase of the experiment by inserting the eavesdropper fly into the same arena with the two demonstrator males that they just observed. We utilized AKASO cameras (EK700) to video record the interaction phase, and then used BORIS, an event logging software, to code the social interactions that occurred between the eavesdropper and the paired males for another 30 minutes (Friard & Gamba, 2016).

On each experimental day, we were able to conduct 8 trials across 2 time blocks: 4 trials began approximately 15 minutes after subjective dawn, and another 4 trials began approximately 1 hour after subjective dawn. Trials were able to be conducted simultaneously through the use of scan sampling (figure 2) during the observation phase, and the use of video recording during the interaction phase, which did not require the presence of a human observer beyond the initial setup. Time block was not found to have a significant effect on male aggression or female mate choice (see Model Covariates in the Supplemental Material). Observers were blind to treatment when using videos to code the social behaviors in the interaction phase, however, the scoring of aggressive behavior during the observation phase was not conducted blind in regards to treatment, as the presence or absence of an eavesdropper fly made this impossible.

### Treatments

We conducted 166 trials over three different experimental treatments, which are explained further below. In 35 trials, we had one or more flies escape after the observation phase, but before or during the interaction phase. When this occurred, we retained the data that we had collected for the observation phase.

#### Female Eavesdropper

In this treatment, we used a female fly as the eavesdropper. For the observation phase of the experiment, a female fly observed an aggressive encounter between the paired males (n=41 trials). For the interaction phase of the experiment, we recorded whether the female mated and, if so, which male the female chose to mate with (n=38 trials).

#### Male Eavesdropper

In this treatment, we used a male fly as the eavesdropper. For the observation phase of the experiment, a male fly observed an aggressive encounter between the paired males (n=43 trials). For the interaction phase of the experiment, we recorded all lunging behavior that the male eavesdropper displayed while interacting with the two males (n=32 trials). We measured only lunging behavior during the second phase of the experiment, as more subtle aggressive behaviors were difficult to confidently assign from our relatively low-resolution videos. We note that lunging behavior is significantly correlated with our four other measures of aggressive behavior in the observation phase of the experiment (see Supplementary Figure 1). Furthermore, measuring only lunging behavior has previously been shown to be an accurate characterization of broad aggressive patterns in male fruit flies (Saltz, 2013).

#### No Eavesdropper/No Social Information

In this treatment, there was no eavesdropper present for the observation phase of the experiment, providing us with a baseline measure of how the paired males would behave without an audience present (n=82 trials). In the interaction phase of the experiment, a male (n=32 trials) or female fly (n=38 trials) who did not observe the aggressive encounter was inserted into the arena. The same behaviors were recorded as described in the male and female eavesdropper treatments, depending on the sex of the fly that was added.

#### Female Eavesdropper Follow Up – No Variation in Paired Male Genotype

After completing our female trials, we observed that there was a strong bias in female mating preferences: in the 59 mating events that occurred, females mated with the genotype 365 male in 51 of those trials. This bias for 365 males obscured any effects of social information on mate choice preferences. As a result, we did a follow-up experiment, where we conducted the female eavesdropper (n=47) and no eavesdropper trials (n=38) using the same protocol as described above, except we used two males from genotype 365 as the paired males engaged in an aggressive encounter. Using this approach, the only difference between the males was their behavior during the observation phase, thus clarifying the effects of social information on female mate choice.

### Ethical Note

No animal ethics approval was required for this study, however, flies were handled with care at all times to minimize stress and damage. We used anesthesia to collect flies, and allowed them to recover for 3 or 4 days before experimental trials. During trials, we handled flies using gentle aspiration. After trials were complete, all flies were euthanized by being placed in a freezer.

## Analysis

The goal of our analysis was to investigate how individuals influenced (observation phase) and used (during the interaction phase) social information. To accomplish this, we created generalized linear mixed models using the glmmTMB package in R (v4.1.0, R Core Team, 2021, Bolker et al., 2009).Generalized linear mixed models allow us to fit a variety of error distributions to our data, include random effects in our models, and account for zero-inflated data (Brooks et al., 2017). We created models that tested how the paired males produced social information, how males used social information, and how females used social information. To investigate how social information altered social decision-making, we quantified which of the paired males performed the most and least aggressive behaviors during the observation phase of the experimental trials. If the paired males displayed the same number of aggressive behaviors, we recorded it as a tie. Our models were then built to identify the factors that influenced male and female responses to the most and least aggressive of the paired males. For all models, we checked model fit by utilizing the DHARMa diagnostic package in R, which conducts simulation-based dispersion tests (Hartig, 2022). We used hypothesis-based model building to include multiple interaction terms between our fixed effects in our models, explained in detail further below. For the final model selection, we only retained interaction terms that improved the fit of the model according to AIC weights.

### Social Information Production Models

Our first model tested how the presence and sex of the eavesdropper influenced the aggressive behavior of the paired males (Social Information Production Model, Table 1). The response variable for this model was the total aggression displayed by the paired males in the observation phase of the experiment. We calculated the total aggression by summing the five measures of aggressive behavior that each fly performed. We had two fixed effects in this model: treatment, which described the presence and sex of an eavesdropper, and age, as we used both 3- and 4-day old flies for all experiments. The date the trial was conducted was included as a random effect to indicate which flies were tested on the same day. A negative binomial error distribution with a logit link function was used for this model, and a term was included to account for zero inflation, as some pairs had no recorded aggressive interactions during the experimental trials. We also created a model to investigate how the eavesdropper’s genotype influenced social information production, this model and its results are reported in the Supplemental Information (Table S1).

**Table 1.** Parameters, test statistics, and their interpretation in the model for social information production. P-values are corrected for multiple testing.

| Parameter | Parameter Estimate [Standard Error] | p-value | Chi-squared value | Interpretation |
| --- | --- | --- | --- | --- |
| Treatment | Female Eavesdropper: -0.18 [0.19]<br>Male Eavesdropper: -0.61 [0.19] | 0.012 | 10.1 | Paired males altered their aggressive behavior based on the presence and sex of an eavesdropper |
| Age | 4 day old flies: 0.41 [0.19] | 0.07 | 4.48 | No effect of age on paired male aggression |
| Date | - | 0.25 | 2.32 | No effect of experimental day on paired male aggression |

### Male Social Information Use Models

Our next set of models tested how access to social information influenced the aggressive behavior and social decision-making of the males that were added into the arena during the interaction phase. We modeled the number of lunges that the added male directed towards the more aggressive paired male (Social Information Use: Male Response to More Aggressive Male Model, Table 2) or the least aggressive paired male (Social Information Use: Male Response to Least Aggressive Male Model, Table 3) during the interaction phase of the experiment.

**Table 2.** Parameters, test statistics, and their interpretation in the model for male social information use towards the more aggressive paired male. P-values are corrected for multiple testing.

| Parameter | Parameter Estimate [Standard Error] | p-value | Chi-squared value | Interpretation |
| --- | --- | --- | --- | --- |
| Treatment | Male Eavesdropper: -:<br>1.51[0.43] | 0.0008 | 12.6 | Access to social information altered how aggressive males were towards previous contest winners |
| Genotype | - | 0.07 | 6.7 | No detected genotypic variation in how males directed their aggression toward the winner |
| Treatment x Genotype | - | 0.0005 | 16.5 | Genotypic variation in how males used social information to direct their aggressive behaviors toward winners |
| Age | 4 day old: -0.18 [0.2] | 0.74 | 0.8 | No effect of age on male aggression |
| Paired Male Aggression | -0.017[0.01] | 0.27 | 2.3 | Measured aggression in the previous contest did not affect how males directed their behaviors toward winners |
| Observer | - | 0.051 | 4.98 |  |
| Date | - | 2 | 0 | No effect of experimental day on aggression |

**Table 3.** Parameters, test statistics, and their interpretation in the model for male social information use towards the less aggressive paired male. P-values are corrected for multiple testing.

| Parameter | Parameter Estimate [Standard Error] | p-value | Chi-squared value | Interpretation |
| --- | --- | --- | --- | --- |
| Treatment | No Social Information – 0.05[0.34] | 1.9 | 0.009 | Access to social information did not alter how aggressive males were towards previous contest losers |
| Genotype | - | 0.05 | 7.5 | Genotypic variation in how males directed their aggression toward the loser |
| Age | 4 day old: -0.52 [0.32] | 0.2 | 2.4 | No effect of age on male aggression |
| Paired Male Aggression | 0.02[0.02] | 0.47 | 1.4 | Measured aggression in the previous contest did not predict how males directed their behaviors toward losers |
| Observer | - | 1.6 | 0.79 |  |
| Date | - | 2 | 0 | No effect of experimental day on aggression |

Fixed effects in the “Social Information Use: Male Response to More Aggressive Male” model included treatment, which described if the male was an eavesdropper or was not allowed to observe social information. We also included a fixed effect for the genotype of the added male, to account for genotypic variation. Our model selection process led us to retain the interaction we tested between genotype and treatment, as this interaction term improved the fit of the model (delta AIC = 1.59). An effect of this term would indicate that genotypes differed in their use of social information. The total aggression displayed by the paired males in the first phase of the experiment was also used as a fixed effect, to determine if our sample of their aggressive behaviors predicted the behavior of the added male. Our final fixed effect was the age of the flies. The date the trial was conducted and the identity of the researcher who coded the trials were included as random effects. We tested an interaction between the aggression of the paired males and treatment, however, that interaction did not improve the AIC of our model (delta AIC = 1.86).

For the “Social Information Use: Male Response to Least Aggressive Male” model, we used the same fixed effects of treatment, genotype, displayed aggression by the paired males, and age, and random effects of date and observer. We also tested for the same interactions as the previous model, however, our model selection process did not support the retention of either the interaction between displayed aggression and treatment (delta AIC = 1.87) or the interaction between genotype and treatment (delta AIC = 3.48). A negative binomial error distribution with a logit link function was used for both male social information use models, and a term was included in both models to account for zero inflation, as some added males had no recorded aggressive interactions during the experimental trials.

### Female Social Information Use Models

Our last models tested how access to social information influenced female mating choices in the interaction phase of the experiment. We only analyzed data from our female follow-up experiment (see above), where females observed and were allowed to interact with two males of their preferred male genotype, 365.

To investigate how females used social information, we first modeled whether females mated, or not (Social Information Use: Female Model, Table 4), with a binary response variable. Females who mated were assigned a value of 1 (n =53 females), and females who did not mate were assigned a 0 (n = 32 females). Treatment was tested as a fixed effect, to investigate how eavesdropping modified the likelihood of female mating. The total aggression displayed by the paired males in the first phase of the experiment was also used as a fixed effect, to determine if aggressive behavior influenced the mating decision of the added female. We included the interaction between treatment and total paired male aggression to investigate how observed social information influenced mating decisions, as it improved model fit according to AIC (delta AIC = 0.51). We also tested the genotype of the added female as a fixed effect, to account for genetic variation. We did not retain the interaction between genotype and treatment, as it did not improve the model’s AIC (delta AIC = 1.46). The date the trial was conducted and the identity of the researcher who coded the trials were included as a random effect.

**Table 4.** Parameters, test statistics, and their interpretation in the model for female likelihood to mate. P-values are corrected for multiple testing.

| Parameter | Parameter Estimate [Standard Error] | p-value | Chi-squared value | Interpretation |
| --- | --- | --- | --- | --- |
| Genotype | - | 0.0027 | 14 | Genotypic variation in likelihood to mate |
| Paired Male Aggression | 0.03 [0.1] | 0.14 | 4 | Measured aggression displayed in the previous contest did not alter mating behaviors |
| Treatment | Female Eavesdropper: 1.63 [0.68] | 0.2 | 3.4 | Access to social information did not significantly predict likelihood to mate |
| Treatment x Paired Male Aggression | - | 0.38 | 2.4 | Observing the aggression of the paired males did not significantly change their likelihood to mate |
| Observer | - | 2 | 0 |  |
| Date | - | 0.81 | 1.2 | No effect of experimental day on aggression |

To further investigate how social information specifically altered female social decision-making, we modeled whether each mated female chose the more aggressive paired male (Social Information Use: Female Response to More Aggressive Male Model) or the least aggressive paired male (Social Information Use: Female Response to Least Aggressive Male Model). We had smaller sample sizes for these models: of the 85 females tested, only 21 mated with the more aggressive male, and only 9 mated with the least aggressive male. These lower samples were a result of many trials resulting in a “tie” between the paired males (n=23 trials), and females often choosing not to mate with either male (n=32 trials). We utilized a binomial error distribution for all female models. The full description of these models and their results can be found in supplemental tables 2 and 3.

### Correction for Multiple Testing

We created multiple models to investigate our hypotheses regarding social information production, use, and social decision-making, which required a correction for multiple testing. For the Social Information Production and Social Information Use: Male Response models, we applied Bonferroni corrections for 2 tests, since we created two models investigating the same dataset for social information production and male social information use. For the Female Response models, we applied Bonferroni corrections for 3 tests. All p-values reported below, in tables 1-4, and in the supplementary tables reflect these corrections.

## Results

### Social Information Production

The paired males altered their behavior based on the presence and sex of the eavesdropper (Figure 3, Social Information Production Model parameter estimates: Female Eavesdropper: −0.18 ± 0.19, Male Eavesdropper: −0.61 ± 0.19, df = 2, p-value = 0.012, Table 1). The paired males were the least aggressive when a male eavesdropper was present (Social Information Production Model: least-squares means for paired male aggression in the male eavesdropper treatment = 1.84). When a female eavesdropper was present, the paired males did not significantly alter their behavior, compared to when there was no eavesdropper present (Social Information Production Model: least-squares means for paired male aggression in the female eavesdropper treatment = 2.27, no eavesdropper treatment = 2.45). Age did not significantly predict paired male aggressive behavior (Social Information Production Model parameter estimate: 4 day old: 0.41 ± 0.19, df = 1, p-value = 0.07, Table 1). We also did not see an effect of eavesdropper genotype on paired male aggression (Supplemental Table 1).

**Figure 3.**
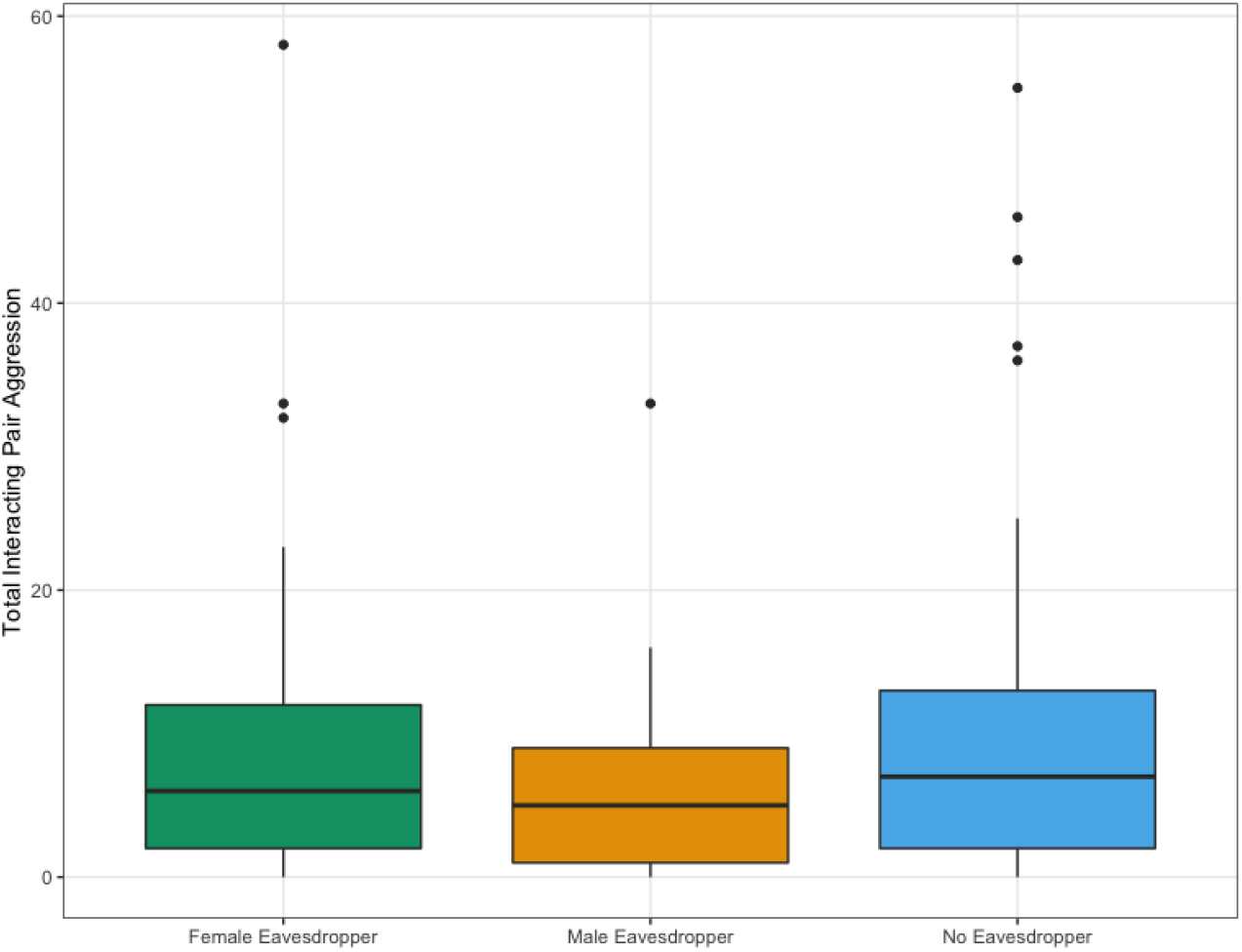
Variation in social information production based on audience identity. The total aggression of the paired males in relation to the sex and presence of an eavesdropper observing them. We calculated total aggression by summarizing five measures of aggressive behavior we recorded for each of the paired males. Analysis via linear mixed models revealed a significant effect of sex and presence of the eavesdropper on paired male aggressive behavior (see text).

### Male Social Information Use

Male flies altered their behavior based on their access to social information. Male eavesdroppers decreased their own aggression towards the more aggressive paired male, compared to males without social information (Social Information Use Male Response to More Aggressive Male Model parameter estimate: Male Eavesdropper: −1.51 ± 0.43, df = 1, p-value = 0.0008, Table 2). We observed genetic variation in how males responded towards the least aggressive paired male (Social Information Use Male Response to Least Aggressive Male Model: X2 = 7.5, df = 2, p-value = 0.047, Table 3), but not the most aggressive paired male (Social Information Use Male Response to More Aggressive Male Model:X2 = 6.74, df = 2, p-value = 0.07, Table 2). We saw a significant genotype by treatment interaction on male aggressive behavior towards the more aggressive paired male. This interaction indicates that genotypes differed in the extent to which they modified their behavior towards the more aggressive paired male based on the ir ability to collect social information during the first phase of the experiment (Social Information Use Male Response to More Aggressive Male Model: X2 = 16.5, df = 2, p-value = 0.0005, Table 2, Figure 4). The total aggression of the paired males in the observation phase of the experiment did not significantly predict how aggressive the male flies would behave towards the more or less aggressive of the paired males (Social Information Use Male Response to More Aggressive Male Model parameter estimate: −0.02 ± 0.01, df = 1, p-value = 0.27; Social Information Use Male Response to Least Aggressive Male Model parameter estimate: 0.02 ± 0.02, df = 1, p-value = 0.47, Table 2 & 3). Age also did not significantly predict male aggressive behavior in any model (Social Information Use Male Response to More Aggressive Male Model parameter estimate: 4 day old: −0.18 ± 0.2, df = 1, p-value = 0.74, Social Information Use Male Response to Least Aggressive Male Model parameter estimate: 4 day old: −0.52 ± 0.32, df = 1, p = 0.2, Table 2 & 3).

**Figure 4.**
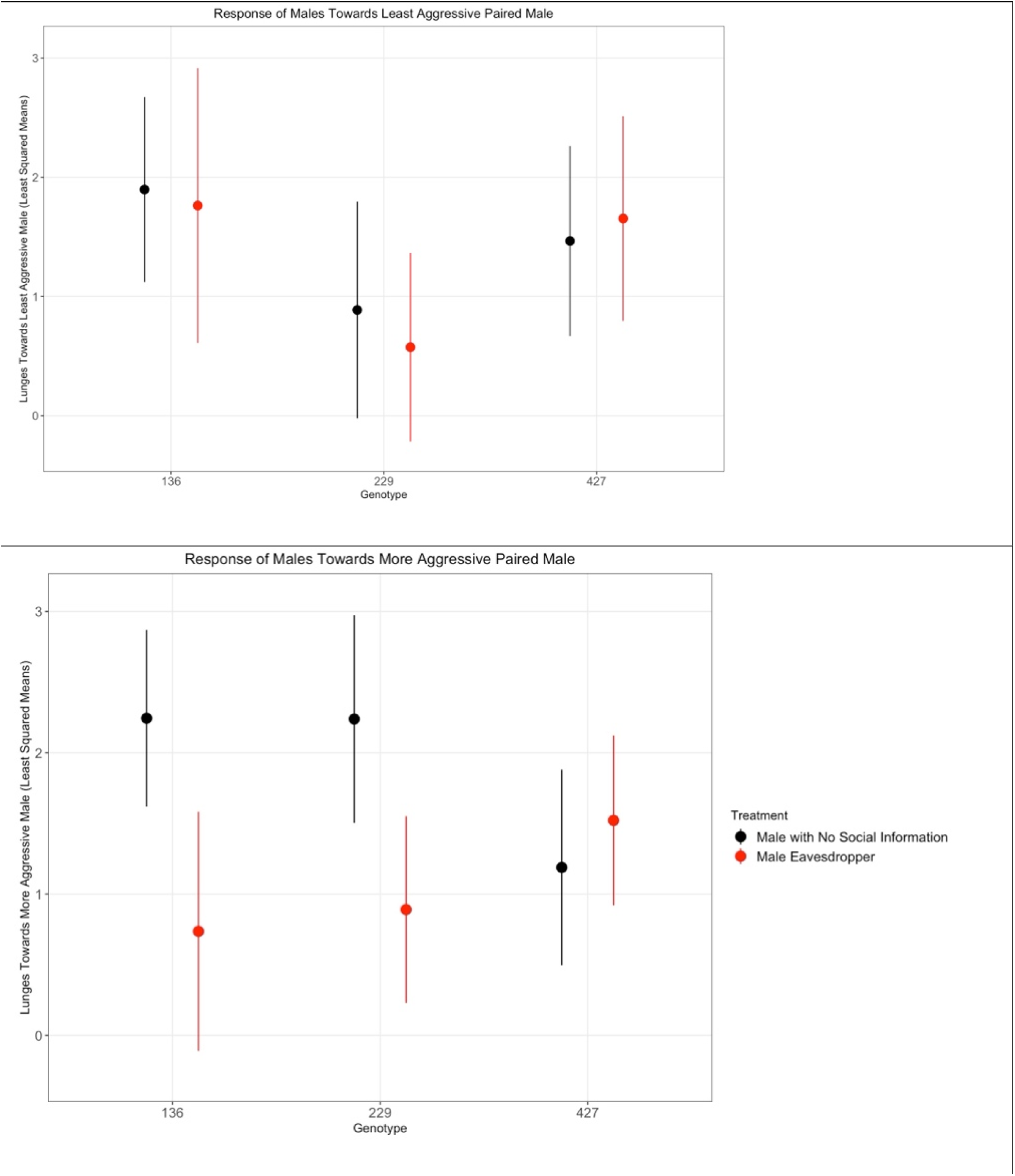
Social decision making in male social information use. The least squared means model outputs for the number of times that the male lunged at the least aggressive or most aggressive of the paired males in the second half of experimental trials is represented on the y-axis in both graphs, and the x-axis in both graphs shows the three tested genotypes. Males with social information lunged significantly less often at observed contest winners than males without social information, but this effect was modulated by genotype.

### Female Social Information Use

Female flies displa yed strong genotypic differences in likelihood to mate (Social Information Use Female Model: X2 = 14.01, df = 2, p-value = 0.003, Table 4). Specifically, females from genotype 229 almost always mated (females mated in 27/29 trials), and females from genotype 136 rarely mated (females mated in 8/27 trials). Neither access to social information nor the displayed aggression of the interacting pair significantly predicted the likelihood of mating (Social Information Use Female Model parameter estimates: treatment Female Eavesdropper: 1.63 ± 0.68, df = 1, p-value = 0.2; displayed aggression 0.03 ± 0.1, df = 1, p-value = 0.14, Table 4). The interaction between total aggression and treatment was also not significant in predicting female mating likelihood (Social Information Use Female Model: X2 = 2.4, df = 1, p-value = 0.38).

None of our fixed or random effects significantly predicted whether the female mated with the more aggressive paired male, or the least aggressive paired male; full results are presented in supplementary tables 2 and 3.

## Discussion

Traditionally, social information use and production have been studied separately, which limits our understanding of how these two processes interact as they occur simultaneously in nature (Bonnie & Earley, 2007; McGregor, 2005). Considering the linkages between social information production and use can improve our predictions about how behaviors spread in populations and identify the causes of individual differences in behavioral strategies (Aplin & Morand-Ferron, 2017; Chimento et al., 2022). However, despite these important implications of the feedback loop on the evolution of social behaviors, we know of no empirical studies of the feedback loop that have been conducted in the context of social eavesdropping.

In this study, we experimentally demonstrate the existence of a feedback loop between social information production and use in an aggressive context in male fruit flies. We found that the presence of a male eavesdropper substantially decreased the aggression of two paired males, supporting an audience effect (Figure 3, Table 1). Furthermore, observing social information caused eavesdropping males to change their aggressive behaviors: male eavesdroppers lunged at the more aggressive paired male less often, compared to males who did not have the opportunity to eavesdrop (Figure 4, Table 2). We also identified genetic variation in male social information use – different genotypes had different responses after observing an aggressive encounter. Overall, these findings indicate that there is a bidirectional transfer of information between the participants and the eavesdroppers in an aggressive context (Figure 5).

**Figure 5.**
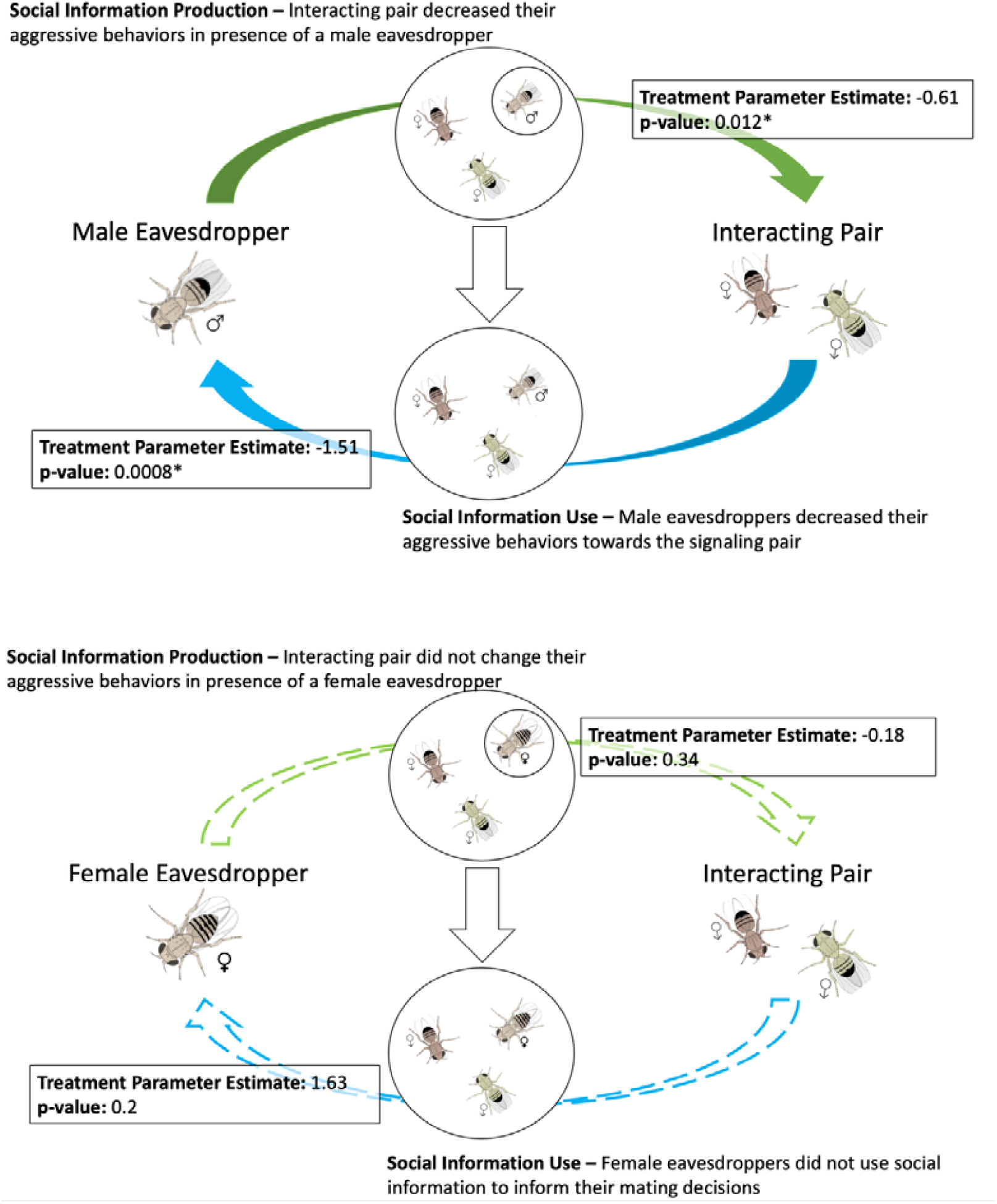
Diagrams depicting how our study’s findings support the existence of a sex-specific feedback loop between social information use and production. Males simultaneously influence the production of and use social information resulting from an aggressive contest. Females do not influence social information production and do not use social information resulting from an aggressive contest.

Social eavesdropping can benefit individuals by allowing them to assess their opponents before engaging in aggressive behavior, thus reducing the costs of fighting (Earley, 2010; Johnstone, 2001). We found a reduction in aggressive behavior from both the paired males when observed by male eavesdroppers, and male eavesdroppers when allowed to interact with the paired males. Thus, the feedback between social information use and production produced substantially lower overall aggression than would be predicted by simply studying pairs of males without eavesdroppers (e.g., in our no eavesdropper treatment), or investigating male eavesdropping without also considering its effect on social information production (e.g., in our no social information treatment). Our results are therefore consistent with previous work that shows obtaining social information about the fighting ability of an opponent may allow individuals to avoid prolonged or escalated encounters (Beltrão et al., 2023; Magnhagen, 2006; Tibbetts et al., 2020); notably, these previous results were obtained without accounting for any influence of the eavesdropper presence on the social information that was produced. Our qualitatively similar findings suggest that, while studying this feedback better informs our understanding of each individual’s behavior, they do not qualitatively alter the patterns (i.e., that individuals use social eavesdropping to avoid potentially costly aggressive encounters) that others have previously observed, at least in this case.

While we are confident that observation of an aggressive encounter altered male eavesdropper behavior, we cannot be certain about the exact social information that the male eavesdroppers were using to inform their decision-making. Winner-loser effects have been documented in flies, providing evidence that flies can learn from aggressive contests – at least, those in which they have directly participated (Penn et al., 2010; Yurkovic et al., 2006). Despite this, our measure of the aggressive behavior of the paired males did not directly influence either male eavesdroppers or males without social information (tables 2 and 3). Eavesdroppers could have extracted information from a variety of sources in the arena, such as observations of how the paired males interacted, chemical signaling behaviors, and/or sounds produced by the paired males (Kent et al., 2008; Versteven et al., 2017).Further research will be necessary to identify mechanistic relationships between eavesdropping, paired male behavior, and the eavesdropper’s later aggression.

Another mechanism that could cause male eavesdroppers to change their behavior in response to the paired males is priming. Priming occurs when observing an aggressive encounter alters an individual’s later behaviors, however, this change in behaviors is expected to be broad and generalize to all future opponents (Clotfelter & Paolino, 2003; Cruz & Oliveira, 2015; Earley et al., 2005; Hsu et al., 2006; Tibbetts et al., 2020). Social eavesdropping, on the other hand, is expected to be individual-specific – the eavesdropper remembers information and learns from the behaviors of those they observed (McGregor, 2005). While we cannot rule out any effects of priming, our results are more consistent with social eavesdropping, as we saw an effect of social information (i.e., treatment) on male behavior that was dependent upon the specific identity of the paired males, i.e., if they were more or less aggressive during the aggressive contest (Table 2). Interestingly, we found that males who did not have the opportunity to eavesdrop on the fight disproportionately attacked the more aggressive of the paired males (Figure 4). With no prior social information to rely on, these males aggressive behaviors were most likely influenced by their personal information and direct experiences once in the arena. For example, fights between the naïve male and the more aggressive paired male may have been instigated by the paired male, leading to increased naïve male aggression as a retaliatory response. In contrast, male eavesdroppers seemingly utilized social information to avoid and engage in fewer aggressive encounters with the more aggressive male. Thus, we find support for a transfer of social information from the paired males to the male eavesdroppers that resulted in a change in behavior that was specific to the individuals that were observed. The detailed cognitive mechanisms underlying this information transfer representing an interesting area for future study. The feedback loop between social information use and production proposes that the presence of a social eavesdropper should cause an adjustment in social information production, which then would lead to changes in eavesdropper behavior. In our study, the paired males displayed similar aggression levels whether a female was observing them or if there was no eavesdropper present (Figure 3). Further, we did not observe any differences in the mating decisions between female eavesdroppers and females without social information (Table 4). Together, these findings further support the idea that changes in eavesdropper behavior are dependent upon modifications in the social information that is produced in their presence.

Genetic variation significantly explained both male and female behaviors in our experiment. Female genotype had a strong effect on mating probability, and paired male genotypes also apparently differed substantially in attractiveness, so much so that we had to repeat the experiment to control for this effect. While eavesdropper genotype did not influence social information production, we interestingly found that male eavesdroppers of different genotypes varied in how social information influenced their response to the more aggressive male. Thus in our feedback loop, we find that all male eavesdroppers had access to similar information, yet male genotypes differed in how they modified their behavioral response to social information. We tested only three genotypes; future experiments that account for genetic diversity are critical for a better assessment of how genetic variation augments or abrogates social information feedbacks (Araya-Ajoy et al., 2020; Cantor et al., 2021).

Our study has a few key limitations to note. All of our flies had no prior social experience before the experimental trials began. While this provided a standardized methodology to test all of our flies, different results may be obtained for individuals that have had prior social interactions with aggression and mating behaviors (Taborsky & Oliveira, 2012). Our female data also presented several challenges: any effect of social information was obscured in our initial set of experiments due to a strong preference for one of the paired male genotypes. In our follow-up trials, our paired 365 males were less aggressive on average than the 765/365 paired males. As a result, we often were unable to assign a more and least aggressive male during the experimental trials. Surprisingly, our females also were not always willing to mate, although they were unmated for their entire adult lives. While we were still able to obtain enough data in our follow-up experiment to investigate the feedback loop, continued work is necessary to better understand how females use social information from male-male aggressive contests, if at all.

This study has revealed several areas of further research necessary to continue to better elucidate the relationship between social information use and production. The sex differences that we observed emphasize the need for continued testing and modeling of these processes in diverse populations, as accounting for individual differences is vital for accurate predictions of social information spread in populations (Duboscq et al., 2016). Additionally, we need a better understanding of the selection pressures in social networks that are introduced via this feedback loop. In this system for example, in order for increased aggression to be adaptive, the benefits must outweigh the costs, but these will likely change depending on the social environment (Duckworth, 2006). Further research can tackle this by quantifying the costs and benefits of plasticity in social information production and use.

Overall, our study lends the first empirical support for a feedback loop between how social information is produced and used in different social contexts. Utilizing the social information feedback framework to investigate animal behaviors can contribute to a better understanding of how behaviors related to conflict, sociality, and group living are expressed and evolve.

## Declarations

## Acknowledgments

We thank Lea Pollack and other members of the Saltz Lab for helpful discussions that improved the manuscript, and we thank Anja Hartge, Keshav Wagle, Erin Vance, Marcelo Dimitri, and Autumn Hildebrand for assistance with fly maintenance and data collection. Funding support for this research was provided by the National Science Foundation (NSF IOS-1856577 and NSF DEB-2217557 to JBS, NSF GRFP-1842494 to MH), and by the Houston Livestock Show & Rodeo First Year Graduate Fellowship (to MH).

## Author Contributions

MH and JBS - Conceptualization; MH - Data collection; MH and JBS - Formal analysis; MH and JBS – Writing original draft, review & editing

## Data Availability Statement

The datasets and code are available on figshare - https://figshare.com/projects/Eavesdropping_and_the_audience_effect_in_fighting_fruit_flies_reveal_feedback_between_social_information_use_and_production/186912

