## Supplemental Material for "Eavesdropping and audience effect in fighting fruit flies reveal feedback between social information use and production"

*Social Information Production Model*

Our follow-up model tested how the genotype of the eavesdropper influenced the aggressive behaviors of the paired males (SI Production Model). Only trials where the eavesdropper was present (male and female eavesdropper trials) were included in this model (n=84). The response variable for this model was the total aggression displayed by the paired males in the first phase of the experiment, and we had three fixed effects in this model: treatment, age, and eavesdropper genotype. Date was included as a random effect. We initially included an interaction term between treatment and genotype, however, this term did not improve the AIC of the model (delta AIC = 2.15) so was removed. A negative binomial error distribution with a logit link function was used for this model, and a term was included to account for zero inflation, as some pairs had no recorded aggressive interactions during the experimental trials.

*Female Response to Winner and Loser Models*

To investigate how social information altered female social decision-making, we modeled whether each mated female chose the winner (SI Use Female Response to Winner Model) or the loser (SI Use Female Response to Loser Model) of the aggressive contest. Fixed effects included treatment, female eavesdropper genotype, and displayed aggression by the paired male, along with a random effect of date. Based on our hypotheses, we tested for an interaction between displayed aggression and treatment, and an interaction term between genotype and treatment in both models. The interaction between displayed aggression and treatment did not improve the model fit for the Winner (delta AIC = 1.93) or the Loser models (delta AIC = 0.11). Including the interaction term between genotype and treatment leads to model convergence issues, likely due to smaller sample sizes (see Analysis).

Supplementary Table 1. Effects, test statistics and interpretation of parameters included in the full model for eavesdropper genetic variation on social information production. P-values are corrected for multiple testing.

| Parameter | p-value | Chi | Interpretation |
|---|---|---|---|
| Treatment | 0.015 | 9.79 | Paired males altered their aggressive behavior based on the sex of an eavesdropper |
| Age | 0.074 | 4.36 | No effect of age on paired male aggression |
| Genotype | 1.08 | 1.24 | No effect of eavesdropper genetic variation on paired male aggression |
| Date | 0.24 | 2.41 | No effect of tested experimental day on the aggression of the paired males |

Supplementary Table 2. Effects, test statistics and interpretation of parameters included in the full model for SI Use Female Winner Model. P-values are corrected for multiple testing.

| Parameter | p-value | Chi | Interpretation |
|---|---|---|---|
| Genotype | 0.6 | 0.2 | No detected genotypic differences in likelihood to mate with winner |
| Paired Male Aggression | 2.09 | 0.7 | Amount of aggression displayed in the previous contest did not influence the females decision to mate with the winner |
| Treatment | 0.98 | 0.33 | Access to social information didn't significantly predict likelihood to mate with winner |

Supplementary Table 3. Effects, test statistics and interpretation of parameters included in the full model for SI Use Female Loser Model.

| Parameter | p-value | Chi | Interpretation |
|---|---|---|---|
| Genotype | 2.29 | 0.76 | No detected genotypic differences in likelihood to mate with winner |
| Paired Male Aggression | 2.139 | 0.38 | Amount of aggression displayed in the previous contest did not influence the females decision to mate with the winner |
| Treatment | 2.13 | 0.71 | Access to social information didn't significantly predict likelihood to mate with winner |

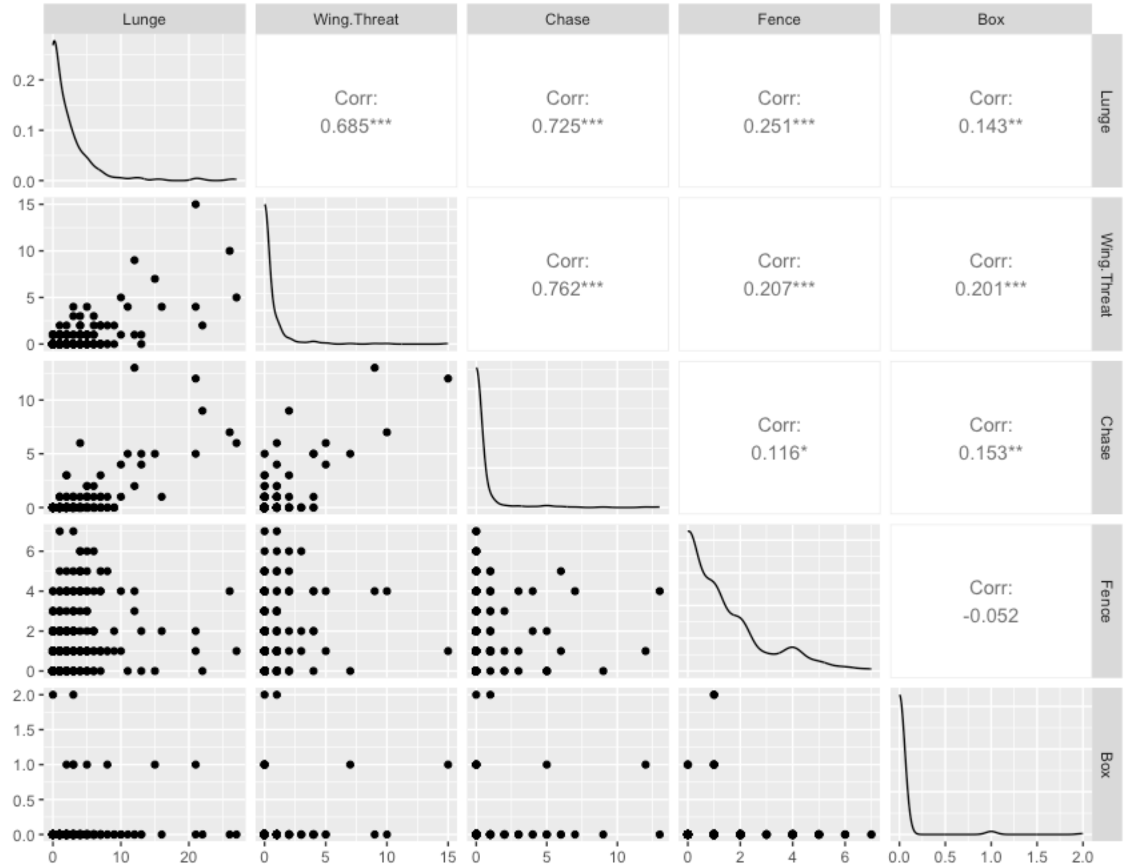


Figure S1. Scatterplot correlation matrix of male aggressive behaviors. The variable distribution of each behavior is on the diagonal, and Pearson correlation is displayed on the right. Male lunging behavior is significantly and positively correlated with the four other measures of aggressive behavior.
